# Alternating handedness motifs in proteins classify structure and cofactor binding

**DOI:** 10.1101/2020.11.17.367490

**Authors:** Shaheer Rizwan, Douglas Pike, Saroj Poudel, Vikas Nanda

## Abstract

Cofactor binding sites in proteins often are composed of favorable interactions of specific cofactors with the sidechains and/or backbone protein fold motifs. In many cases these motifs contain left-handed conformations which enable tight turns of the backbone that present backbone amide protons in direct interactions with cofactors termed ‘cationic nests’. Here, we defined alternating handedness of secondary structure as a search constraint within the PDB to systematically identify these cofactor binding nests. We identify unique alternating handedness structural motifs which are specific to the cofactors they bind. These motifs can guide the design of engineered folds that utilize specific cofactors and also enable us to gain a deeper insight into the evolution of the structure of cofactor binding sites.

## Introduction

Peptide macrocycles, or cyclic peptides require left-handed conformations in secondary structure to accommodate the small radius of curvature.^1^ However, due to the chiral nature of amino acids (with L and D enantiomers), and the exclusivity of the ribosome to synthesize L-amino acids, right-handed conformations are more energetically accessible to protein conformational space and are thus more highly occupied in the protein structure database (PDB).^2^ Glycine, however, is an achiral amino acid and enables most of these left-handed conformations that are present in protein secondary structure.^3,4^ This secondary structure was first characterized by Ramachandran et. al, wherein, they determined that it could be systematically defined by a combination of two angles for each residue, termed phi and psi, which are measured along the protein backbone. ^5^ They calculated the acceptable conformations of protein secondary structure by estimating the favorable and unfavorable interaction of atoms in proteins. Often, it is left-handed conformations that contain glycine and occasionally other amino acids that enable the protein backbone to contort into the low radius of curvature needed for cyclic peptides.^1^ Similar peptides can form the cationic nest as defined by White et. al which is capable of binding metals and other cofactors that are relevant to the origins of life.^6–8^ These cationic nests have direct or indirect backbone amide interactions with the bound cofactor and confer activity or stability to that binding site. Here, we seek to quantitatively analyze a broader space of “nests” that includes multiple nest types (both cationic and anionic, and another type) through a more general approach involving clustering phi/psi angles of contiguous residues that contain alternating handedness conformations

## Methods

### Compiling protein structures

Non-redundant protein structures were downloaded from PISCES (protein sequence culling server) of the Dunbrack Lab with these specifications: sequence percentage identity was <= 30%, resolution <= 2.5 angstroms, R factor = 0.3, sequence length > 40.^9^ Biopython’s PDB module (Bio.PDB) was used to locally download PDB format (i.e.,‘.pdb’) files from the list of non-redundant protein structure, and appropriate chains were selected.^10,11^

### Calculating phi and psi angles

Dihedral angles of phi and psi for every residue were calculated using Bio.PDB and were compiled in individual text files. Each protein’s phi-psi space was then scanned for stretches of alternating positive/negative phi angles. Positive phi indicates a left-handed conformation while negative phi indicates a right-handed conformation. This narrowed our dataset to only regions of proteins that had alternating positive/negative phi angles (alternating handedness). The names and residues of alternating phi/psi regions were compiled into a spreadsheet.

### Calculating backbone angles

Protein backbone curvatures were calculated as the angle formed between alpha carbons (α) (i.e., Cα1–Cα3–Cα5) with a length of five.

### Clustering phi and psi angles

The alternating phi/psi angles were clustered with k-means clustering via scikit-learn.^12^ Clustering was done on the sine and cosine of phi and psi, which created clusters based on backbone structure. Heteroatoms in the vicinity of the backbone were identified (heteroatoms closer than five angstroms from the backbone nitrogen of each residue). Heteroatoms that were part of water molecules were excluded from this data collection due to the number of water molecules that appeared in this analysis.

### Binning conformations

Phi/psi’s were binned into the conformational spaces they fit into within the Ramachandran plot: left/right poly-proline, left/right β-sheet, left/right alpha helix, and another region for phi/psi pairings that did not fit into any of these regions. For example, a 5-residue long sequence of an alternating-handedness alpha helix conformation would be denoted as αR αL αR αL αR (**Fig. 1**). For each cluster, the conformational sequences of all peptides in that cluster were used to create a single consensus sequence for each backbone motif using the biopython’s motif function via objects consensus tool.

**Figure 1).**
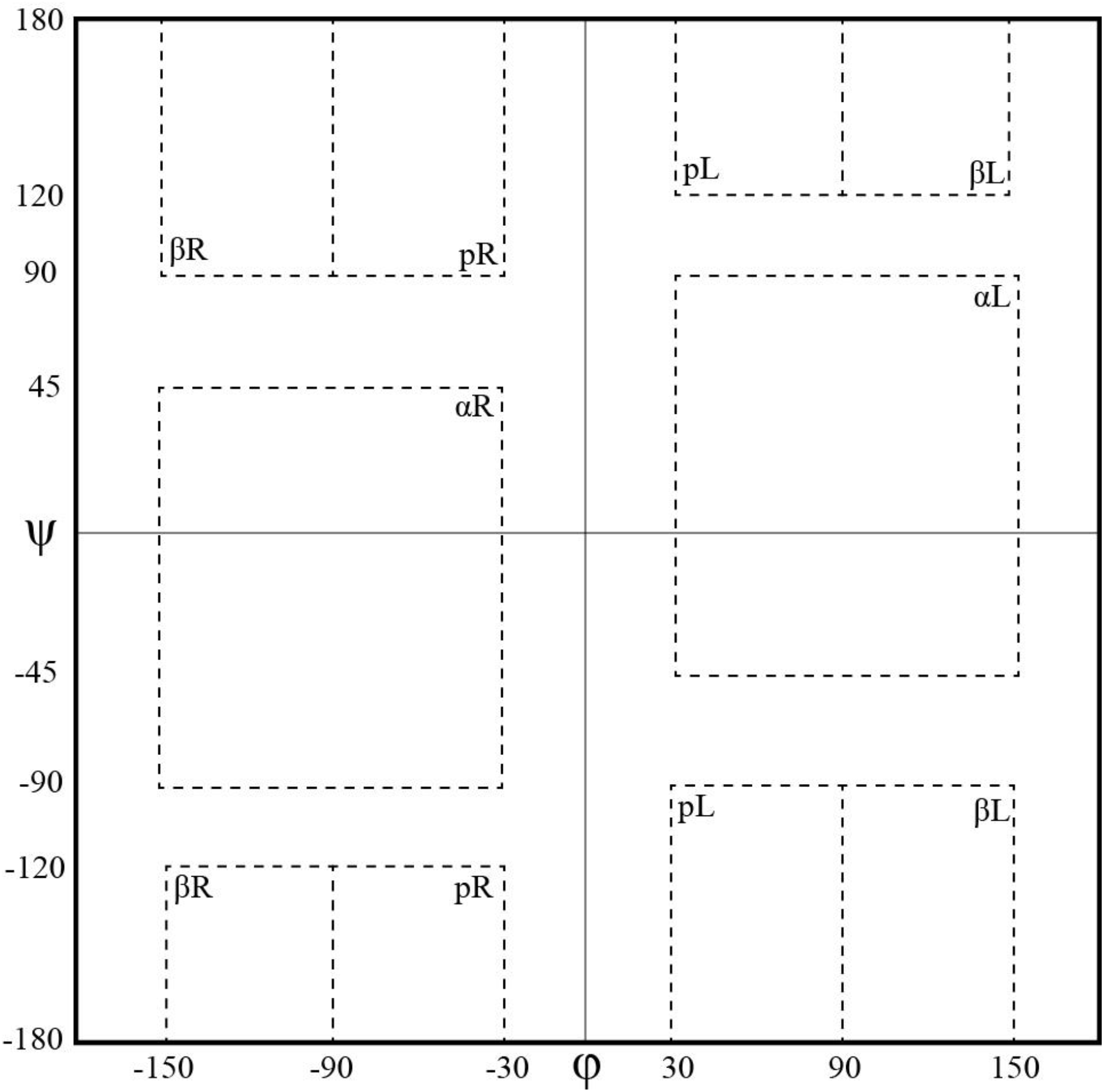
Binning of Ramachandran space - Phi/psi angles were calculated for all proteins; they were binned as seen in the figure. Left-handed conformations have a phi > 0, and right-handed conformations have a phi < 0. Beta (β), polyproline (p) and alpha (α) regions, combined with left/right regions created a conformational ‘alphabet’, used for a written motif. Boundaries for each bin: **αR**: −30 < phi < −150, 45 < psi < −90; **αL**: 30 < phi < 150, −45 < psi < 90; **βR**: −90 < phi < −150, 90 < psi < 180, −180 < psi < −120; **βL**: 90 < phi < 150, −180 < psi < −90, 120 < psi < 180; **pR**: −90 < phi < −30, −180 < psi < 120, 90 < psi < 180; **pL**: 30 < phi < 90, −180 < psi < −90, 120 < psi < 180.

### Graphing and Subclustering

Supplementary graphs of phi/psi of each cluster were also made with matplotlib/seaborn.^13^ This provided a visual overview of the conformational space of each cluster. Clusters that bound a high percentage of a specific cofactor were further resolved by “subclustering” – using the same k-means clustering algorithm on individual clusters. This created subclusters that generally divided clusters by cofactor, creating a cluster binding group and an apo (i.e., without any bound cofactor) group. Representative structures of each cluster were chosen based on the presence of a cofactor and/or the pairwise RMSD of the conformation relative to every member of the cluster.

### Calculating Energies of Theoretical Conformational Space

To see how this distribution of alternating handedness seen in nature compares to a theoretically attainable set of backbones of alternating handedness in terms of energy constraints, we used protCAD to calculate electrostatic and Van der Waals energies of a 7 residue peptide featuring alternating alanine (in right handed conformations) and glycine (in left handed conformations).^14^ This ala/gly heptapeptide was cycled through all possible combinations (a total of 243) of conformational sequences (αR αL αR αL αR, then αR αL αR αL βR, etc.). The specific phi/psi values for each conformation was calculated as the average of all of the phi/psi values of that specific conformation that we initially found in our search in nature. Theoretical conformations had their curvature angles calculated, and this was graphed on a histogram weighted by the energy of the conformation. To do this, the conformations were fitted to a Boltzmann distribution at T = 1500K.

## Results

### Distribution of phi and psi angles

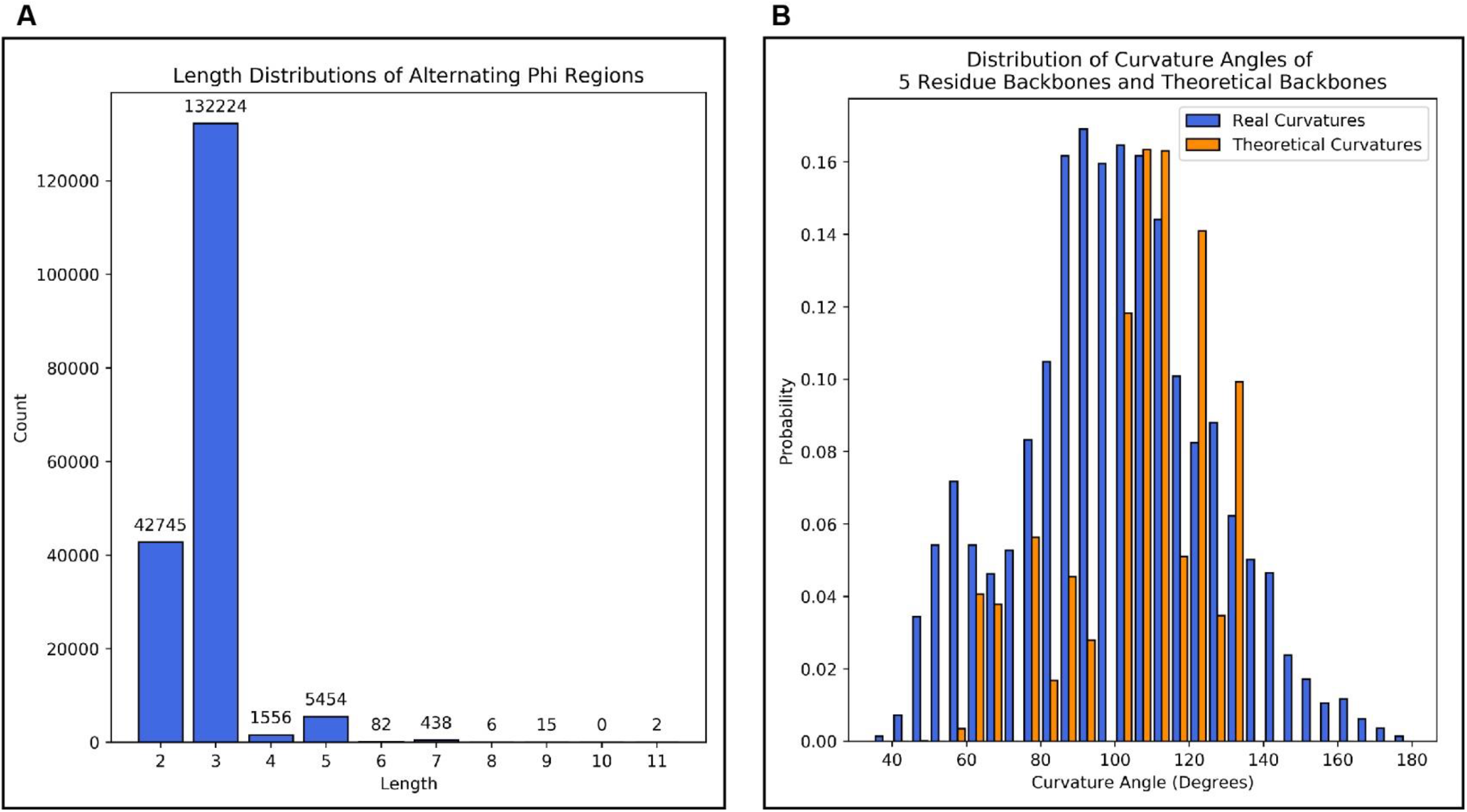

In searching for a pattern of alternating handedness and clustering our motifs based only on phi and psi, we have identified 5 groups of motifs that are not only structurally similar, but also bind similar cofactors, with similar backbone angles, and oftentimes with similar residues. The ability of our approach to identify motifs with these similar characteristics through an analysis of only two parameters, phi and psi, demonstrates the importance of nests of alternating handedness in binding biologically relevant cofactors. Our analysis initially identified a total of 14,376 protein structures from our non-redundant dataset deposited in the PDB, corresponding to a total of 7551 backbones with alternating phi angles. While most were of length five, there were a large number with length seven, and two backbones with length 11 (**Fig. 2A**). We focus on the most abundant group of motifs, those of length 5, which is the shortest peptide for which a β angle can be calculated. The curvature angles in alternating left/right backbones (of length 5) range from ~35 to ~180 degrees, and it shows one broad concentration of angles between 95 and 120 degrees (**Fig. 2B**). Comparing this curvature angle distribution to the theoretically attainable set of conformations in **Fig. 2B**, we observe a similar sort of distribution, with the highest concentration of angles around 110 degrees. There is a relative underrepresentation of theoretical angles between ~75 to 100 degrees. This difference is could be due to the presence of counterion/metal coordination in nests that stabilize the smaller curvature angles in the real set of backbones, which is not represented in the theoretical set.

Our initial clustering approach yielded a total of 25 clusters (**Table S1**). 25 clusters was the maximum number of clusters in which a unique conformational sequence was determined. We subsequently selected four of these clusters that bound a high percentage of a specific cofactor to be ‘subclustered’ and split again. While this was a structural clustering approach, subclustering split these clusters into a holoprotein group and an apoprotein group in all cases.

We identified two groups that resemble cationic nests, two that resemble anionic nests, as well as an additional group which doesn’t that has some SAM/SAH binding potential. For all groups, the sequence logos which display the primary amino acid sequence highlight the importance of glycine in positions 2 and 4, needed to accommodate the left-handed conformation. Also, comparing the Ramachandran distribution between clusters makes some trends clear. For example, comparing group 23 to group 3, the substitution of a polyproline region to the N terminus of group 23 (**Fig. S1A-B**) results in a tightening up of the curvature angle, altering it from ~120/100 degrees to ~90/85 degrees. Similarly, a beta conformation at the C terminus also tightens up the curvature angle relative to pure alpha left/right, going from ~120/100 degrees in 23 to ~100/70 degrees in 5 (**Fig. S1B-C**).

## Cationic Nest

We identify two cationic nests which are Ferredoxin and Rossman-like motifs with amides pointing towards the cofactor which are thought to increase cofactor binding specificity and stability.

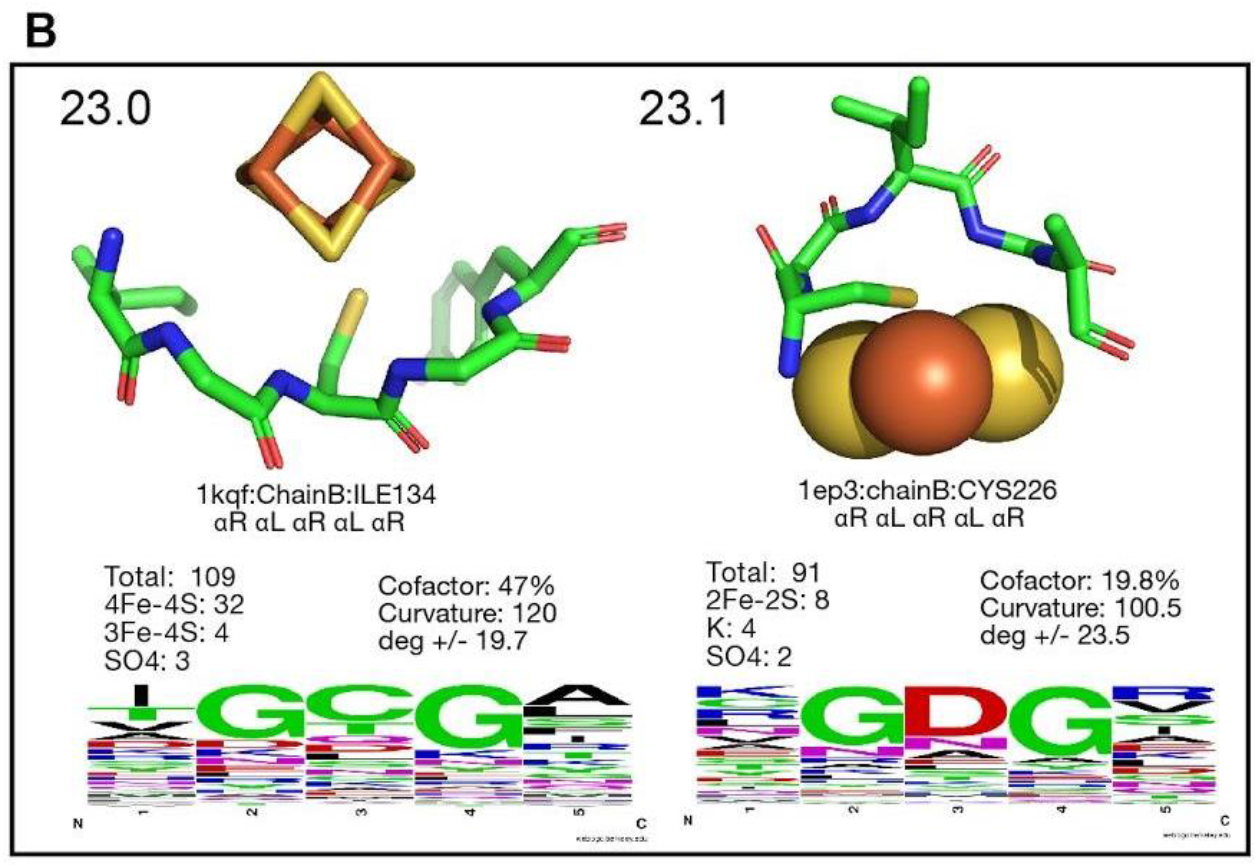

### 1) Ferredoxin

The ferredoxin fold identified in Figure 3B is a highly conserved motif capable of electron transfer by binding a 4Fe-4S cluster in its center.^15–17^ Ligand-metal interactions between cysteine and the metal cluster help to stabilize the cluster in addition to amide proton interactions with sulfurs in the cluster.^18^ Ferredoxin has played an important role in prebiotic proteins since it is one of ten super-folds that has given rise to the diverse landscape of protein structure through evolutionary time.^19^ While the extant ferredoxin does not exhibit heterochirality, there is potential for a prebiotic heterochiral ferredoxin that might have existed on early Earth, considering that a recent heterochiral ferredoxin mimic was designed, demonstrating stability over many redox cycles.^18^ This represents a potential class of proteins that might have existed during early earth conditions, in which D-amino acids have been used in the left handed backbone conformation positions to stabilize the alternating conformation.^18^ Group 23.0 (**Fig. 3B**) represents the ferredoxin fold binding a 4Fe-4S cluster. It features a very similar average curvature to known ferredoxin folds. As expected, a central cysteine signal is involved in iron binding. The apo group (Group 23.1) has a smaller curvature (~100 degrees) and also can bind 2Fe-2S clusters. An analysis of the backbones that bind 2Fe-2S clusters reveals that it requires a smaller curvature angle relative to the group (~75 degrees), and also binds the cluster through a cysteine interaction at the first position. Additionally, group 23 has 2 ‘shells’ of charge - a positively charged inner shell assisting in FeS binding from amide nitrogens, and a negatively charged outer shell from backbone carbonyls.

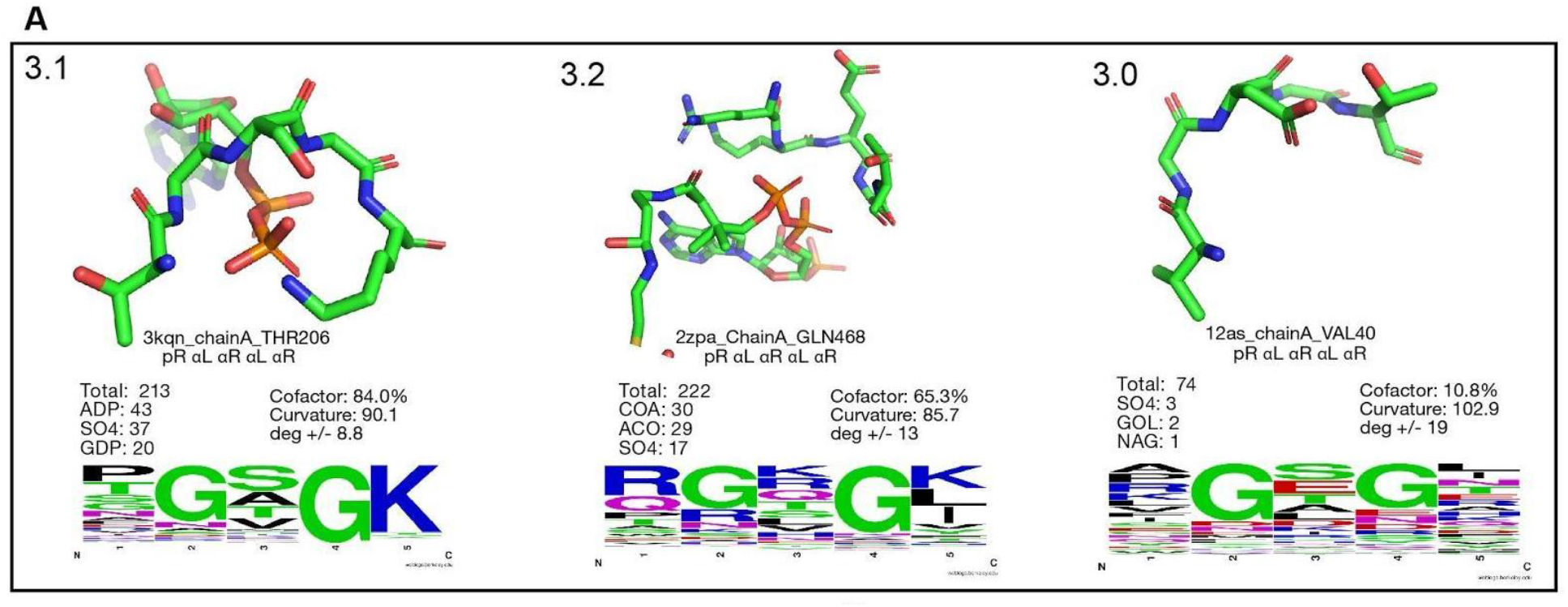

### 2) Rossman

The Rossmann fold is made of alternating beta strand and alpha helical segments, with a highly conserved initial βαβ structure.^20^ This specific portion binds the ADP portion of NADP, FAD, etc.^19^ With its relevance to metabolism, the Rossmann fold has an important evolutionary role.^21,22^ Its cofactor binding is similar to that of group 3.1 (**Fig. 3A**). Here, the strong lysine signal at the fifth residue seems to be critical for ADP binding, through lysine side chain hydrogen bonding with oxygen of phosphate groups. Group 3.1 also looks similar to the C-terminus of the Walker A motif.^23^ In group 3.1, 84 percent binds heteroatoms: mostly ADP, SO4, and GDP. Group 3.2 has a similar curvature to group 3.1, but it primarily binds coenzyme A and acetyl coenzyme A, also around the diphosphate group. Positively charged residues are used in right-handed positions. Group 3.0, the apo group holds a similar occupation in Ramachandran space but features a larger curvature angle and no real selectivity for the particular residues that dictate diphosphate binding.

## Anionic nest

We also extend the idea of a ‘nest’ to also include 2 anionic nests that we identified in our analysis, binding through either side chain interactions or backbone carbonyls.

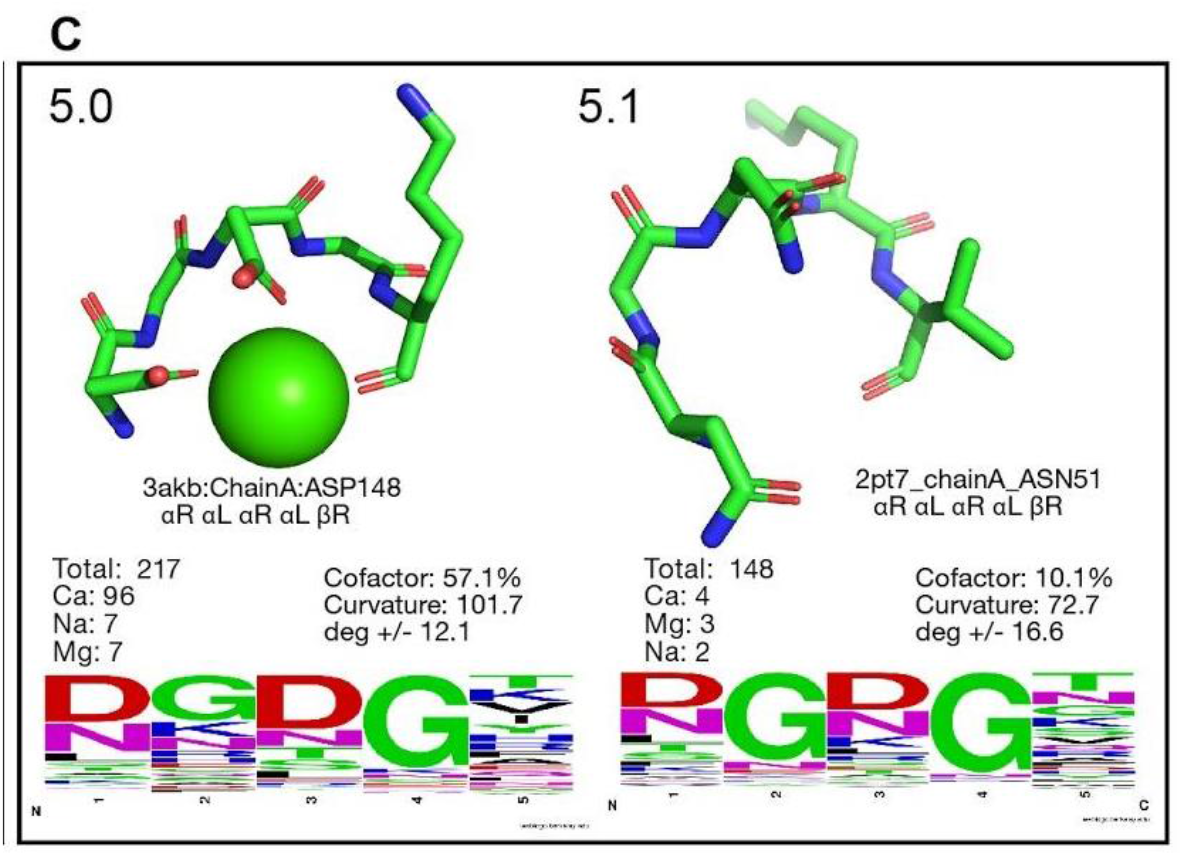

### 1) Side chain Anionic Nest

Group 5.0 also binds calcium (Ca^2+^) but through side chain interactions primarily with aspartates at the first and third positions (**Fig. 3C**). The beta right conformation at the fifth position allows for additional stabilization through backbone carbonyl interactions. The apo group, 5.1, cannot bind Ca due to its smaller curvature angle. Only 5.0 has a sufficiently large curvature angle (~100 degrees) for binding. Group 5 can also be thought of as hosting 3 ‘shells’ of charge, an inner anionic shell from side chain interactions, a middle cationic shell from amide nitrogens, and a third outer anionic shell from backbone carbonyls. This sort of concentric charge distribution is highlighted in **Fig. S2A**.

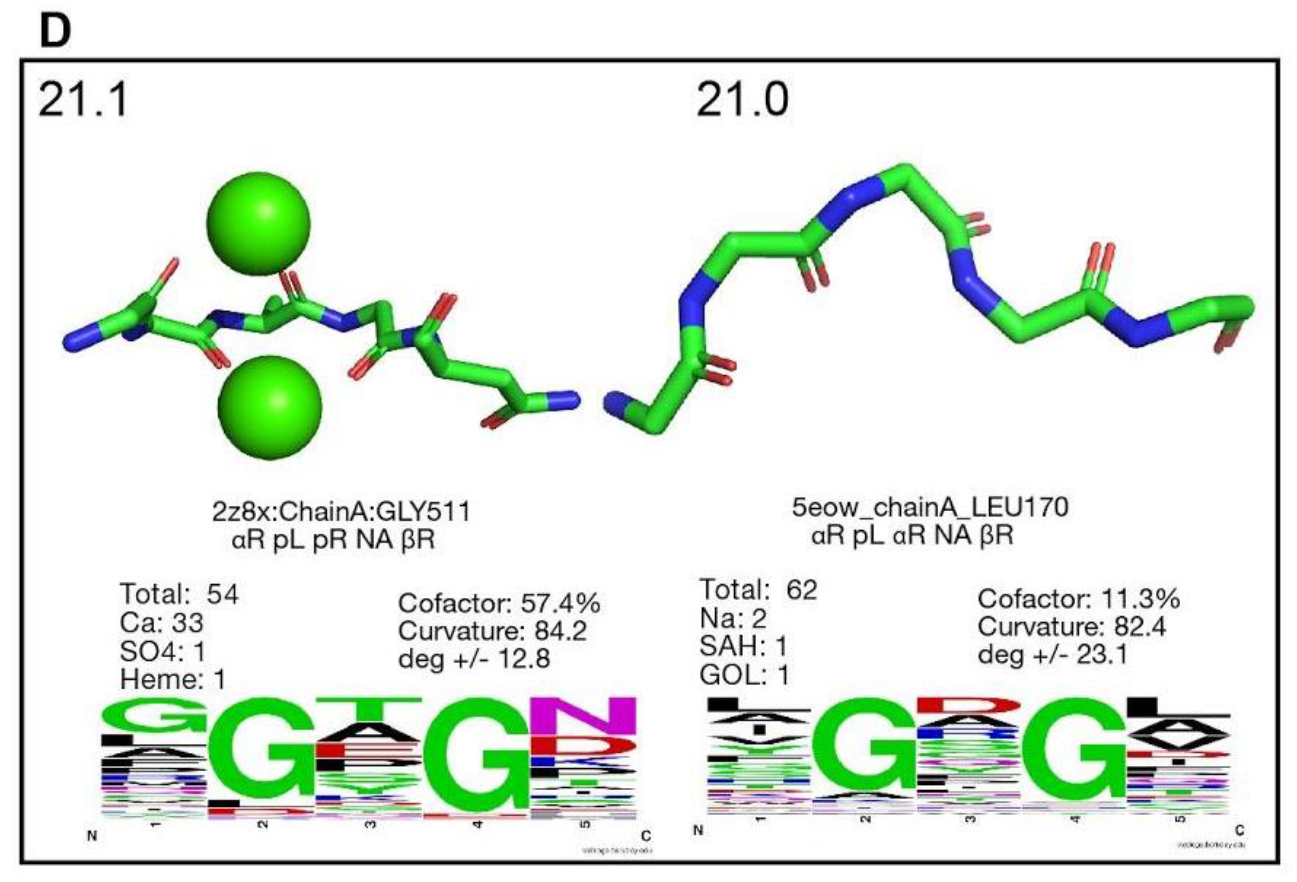

### 2) Backbone Anionic Nest

Extending the idea of the more common cationic nest, in which the amide protons oftentimes form direct electrostatics interactions with the cofactor, anionic nests typically coordinate cations and interact with carbonyls in the backbone of the amino acids in the nest.^7^ As shown in Fig. 3D calcium ions can be found in these anionic nests and they can bind multiple cations per protein, as in PDB id: 2zbx.^24^ The alternating polyproline motif properly aligns the backbone carbonyls to create this anionic nest, which forms a negatively charged shell surrounding the positive cation. The outermost shell consists of another cationic ring from amide nitrogens (**Fig. S2B**).

An additional interesting element is how cofactor binding is dictated by specific and tightly grouped regions in the Ramachandran plot. This is seen especially in cofactor binding groups 3.1 and 3.2, whose alpha-right region has a notable divide between residues 3 and 5 (**Fig. S1A**). This contrasts with the alpha-right region of 3.0, the apo group, where residues 3 and 5 are entirely mixed throughout the alpha-right bin. The same phenomenon is seen group 23, where the phi/psi distribution of residues 1, 3, and 5 in 23.0 are tighter and less homogeneous compared to the apo group, 23.1 (**Fig. S1B**). A similar situation exists in group 5.0. Here, however, alpha-left residues (residues 2 and 4) and beta right residues (residue 5) are grouped distinctly and tightly in 5.0, whereas alpha-right residues (1 and 3) are intermingled with each other within that bin (**Fig. S1C**). The opposite situation is seen in the apo group (5.1), where there is an interesting separation in the phi/psi occupancy of residues 1 and 3. It is interesting to note how this distinct occupancy of residues 1 and 3 in phi/psi space do not lead to any specific cofactor binding, despite the opposite being true for groups 3 and 23. This phenomena is not seen in group 21, however.

## Other

Outside of the realm of cationic/anionic nests exists a space of intriguing conformations that do not clearly resemble either of the two. Here, we report 2 structurally distinct motifs that show potential for SAM/SAH binding, with both motifs seeming having a more extended conformation, unlike the nests described above.

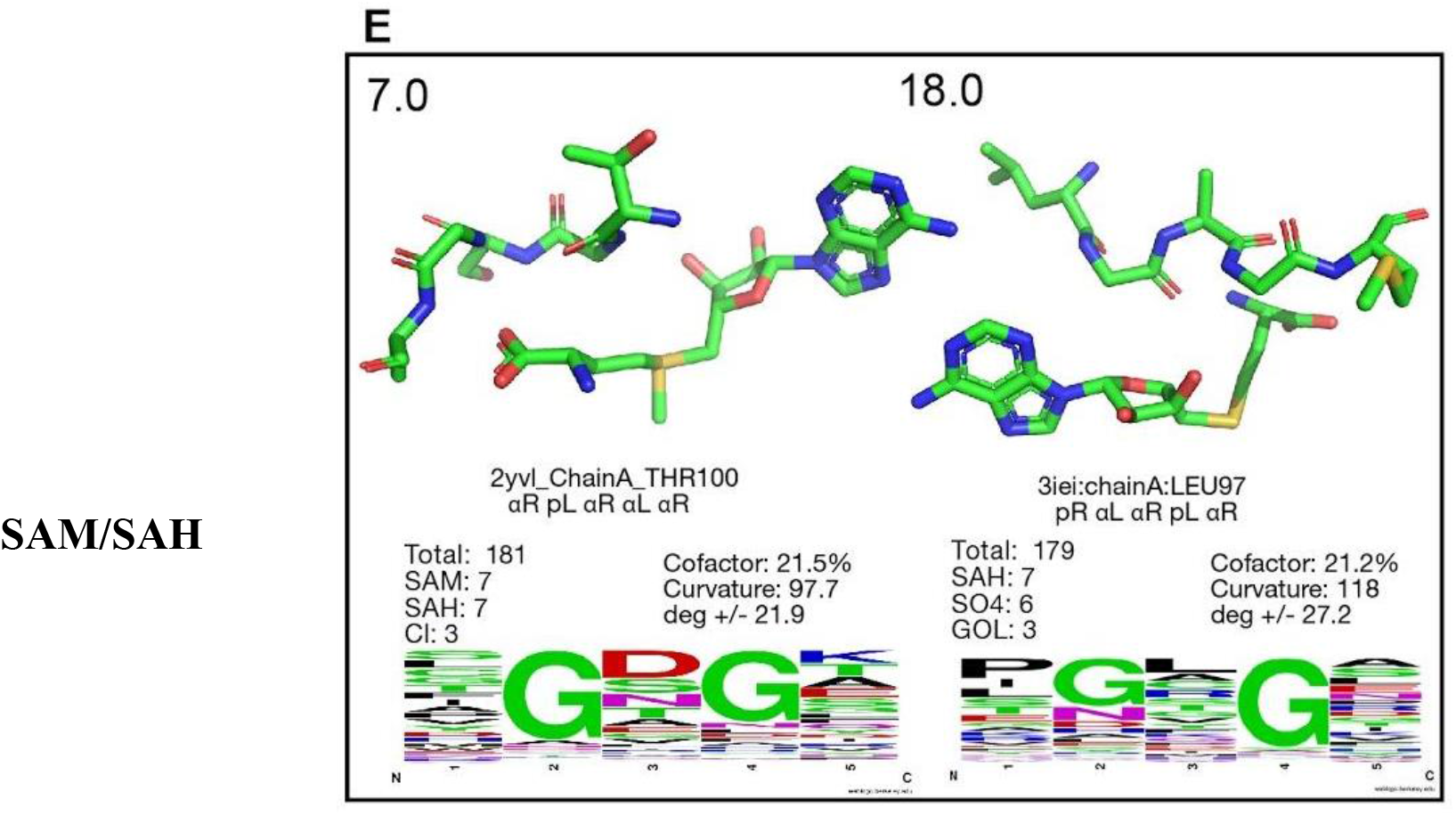

In group 7, s-adenosylmethionine (SAM) is seen, which is a commonly used substrate to transfer methyl, alkyl and adenosyl groups in various metabolic reactions.^25^ Widely found in all three domains of life (i.e., archaea, bacteria and eukarya), SAM-dependent enzymes have been hypothesized to have been present in our earliest ancestors commonly known as the Last Universal Common Ancestors (LUCA) partly because of its involvement in tRNA modification.^26^ However, since its discovery in 2001, it has been found to be involved in several other crucial metabolism, biogenesis and modifications.^27^ Importantly, they are also involved in maturation of complex metalloenzymes that includes the maturation of the active site (i.e., H-cluster) of iron-iron (FeFe) hydrogenase, Fe-only hydrogenase and catalytic iron-molybdenum (FeMo) cofactor of nitrogenase.^25^

Demethylation of SAM creates another cosubstrate, S-Adenosyl-L-homocysteine (SAH) which is involved in biosynthesis of cysteine and vitamin B6.^28^ Both SAM and SAH are bound by an interaction between the carboxylate end of SAM/SAH. 18 (Fig. 3E) features a similar mode of binding but with a larger curvature.

## Discussion

It is clear that these alternating handedness conformations are consistent and specific to cofactors across the PDB, where various radii of curvatures are geometrically and electrostatically matched to the bound cofactor. In each phi/psi clustered group there is significant specificity of interaction for a given subset of cofactors and in many cases predominantly one cofactor. Likely this is due to the favorable electrostatic interactions coming from backbone amide protons or carbonyl oxygens that can be oriented in a charged nest around the cofactor, where these interactions can stabilize cofactor binding in spite of sidechain ionization or other related reactions that may be essential for the functioning of the site. The consistency of the conformational specificity of the structural sequence for each cofactor likely implies that these conformations were selected over the course of evolution for their ability to aid in binding while sidechain chemistry can be diversified for various functions utilizing the redox potential of a given cofactor.

In defining these structural motifs in proteins, we further elucidate the core principles that guide specific catalyst binding and how these structures can translate to function. In that way, these alternating handedness motifs provide some direction for protein design including heterochiral protein design, which can utilize D-amino acids to stabilize the left-handed conformations in order to prearrange the structure for cofactor binding stability and specificity. The glycine residues at positions 2 and 4 can be replaced with D-amino acids, which can easily accommodate the left-handed conformation. Our results also provide direction on designing inverso cyclic peptides. The entire motif can be stereochemically inverted, with glycine replaced by L amino acids and the other positions replaced by D-amino acids. Considering the radius of curvature of some of these motifs, they can be extended out to form a macrocycle with side chain specificity and specific cofactor binding. Our identified motifs can also play a role within designs that incorporate helix caps with glycines in αL conformations, or in heterochiral designs where D-amino acids with specific side chains operate as C-terminal caps. Such capping motifs can allow the chain to maintain stabilizing interactions across the entire helix and potentially increase thermodynamic stability.^29,30^ Apart from demonstrating how specific patterns of alternating handedness motifs can specify cofactor binding, these motifs also give rise to various avenues of protein design that can leverage this unique pattern of left and right-handed conformations.

## References

1. Hosseinzadeh, P. et al. Comprehensive computational design of ordered peptide macrocycles. Science (80-.). 358, 1461–1466 (2017).

2. Englander, M. T. et al. The ribosome can discriminate the chirality of amino acids within its peptidyl-transferase center. Proc. Natl. Acad. Sci. U. S. A. 112, 6038–6043 (2015).

3. Kimura, S., Kanaya, S., Ishikawa, K., Morikawa, K. & Nakamura, H. Glycine or Asparagine at the left-handed conformation enhances protein thermostability. Protein Eng. Des. Sel. 6, 17–17 (1993).

4. Nicholson, H., Söderlind, E., Tronrud, D. E. & Matthews, B. W. Contributions of left-handed helical residues to the structure and stability of bacteriophage T4 lysozyme. J. Mol. Biol. 210, 181–193 (1989).

5. Ramachandran, G. N., Ramakrishnan, C. & Sasisekharan, V. Stereochemistry of polypeptide chain configurations. Journal of Molecular Biology vol. 7 95–99 (1963).

6. Milner-White, E. J., Nissink, J. W. M., Allen, F. H. & Duddy, W. J. Recurring main-chain anion-binding motifs in short polypeptides: Nests. Acta Crystallogr. Sect. D Biol. Crystallogr. 60, 1935–1942 (2004).

7. Watson, J. D. & Milner-White, E. J. A novel main-chain anion-binding site in proteins: The nest. A particular combination of φ,ψ values in successive residues gives rise to anion-binding sites that occur commonly and are found often at functionally important regions. J. Mol. Biol. (2002) doi:10.1006/jmbi.2001.5227.

8. James Milner-White, E. Protein three-dimensional structures at the origin of life. Interface Focus vol. 9 20190057 (2019).

9. Wang, G. & Dunbrack, R. L. PISCES: A protein sequence culling server. Bioinformatics 19, 1589–1591 (2003).

10. Hamelryck, T. & Manderick, B. PDB file parser and structure class implemented in Python. Bioinformatics (2003) doi:10.1093/bioinformatics/btg299.

11. Mckinney, W. Data Structures for Statistical Computing in Python. PROC. 9th PYTHON Sci. CONF 51 (2010).

12. Pedregosa Fabian et al. Scikit-learn: Machine Learning in Python Gaël Varoquaux Bertrand Thirion Vincent Dubourg Alexandre Passos PEDREGOSA, VAROQUAUX, GRAMFORT ET AL. Matthieu Perrot. J. Mach. Learn. Res. 12, 2825–2830 (2011).

13. Hunter, J. D. Matplotlib: A 2D graphics environment. Comput. Sci. Eng. 9, 99–104 (2007).

14. Pike, D. H. & Nanda, V. Empirical estimation of local dielectric constants: Toward atomistic design of collagen mimetic peptides. Biopolymers 104, 360–370 (2015).

15. Chandonia, J. M., Fox, N. K. & Brenner, S. E. SCOPe: Manual Curation and Artifact Removal in the Structural Classification of Proteins – extended Database. J. Mol. Biol. 429, 348–355 (2017).

16. Raanan, H., Pike, D. H., Moore, E. K., Falkowski, P. G. & Nanda, V. Modular origins of biological electron transfer chains. Proc. Natl. Acad. Sci. U. S. A. 115, 1280–1285 (2018).

17. Adman, E. T., Sieker, L. C. & Jensen, L. H. Structure of a bacterial ferredoxin. J. Biol. Chem. 248, 3987–96 (1973).

18. Dongun Kim, J. et al. Minimal heterochiral de novo designed 4Fe-4S binding peptide capable of robust electron transfer. J. Am. Chem. Soc. (2018).

19. Raanan, H., Poudel, S., Pike, D. H., Nanda, V. & Falkowski, P. G. Small protein folds at the root of an ancient metabolic network. Proc. Natl. Acad. Sci. 117, 7193–7199 (2020).

20. Hanukoglu, I. Proteopedia: Rossmann fold: A beta-alpha-beta fold at dinucleotide binding sites. Biochem. Mol. Biol. Educ. 43, 206–209 (2015).

21. Rao, S. T. & Rossmann, M. G. Comparison of super-secondary structures in proteins. J. Mol. Biol. 76, 241–256 (1973).

22. Hanukoglu, I. Conservation of the Enzyme–Coenzyme Interfaces in FAD and NADP Binding Adrenodoxin Reductase—A Ubiquitous Enzyme. J. Mol. Evol. 85, 205–218 (2017).

23. Walker, J. E., Saraste, M., Runswick, M. J. & Gay, N. J. Distantly related sequences in the alpha- and beta-subunits of ATP synthase, myosin, kinases and other ATP-requiring enzymes and a common nucleotide binding fold. EMBO J. 1, 945–951 (1982).

24. Sugimoto, H. et al. Crystal Structure of CYP105A1 (P450SU-1) in Complex with 1α,25-Dihydroxyvitamin D 3 ^†, ‡^. Biochemistry 47, 4017–4027 (2008).

25. Broderick, J. B., Duffus, B. R., Duschene, K. S. & Shepard, E. M. Radical S-adenosylmethionine enzymes. Chemical Reviews vol. 114 4229–4317 (2014).

26. Weiss, M. C., Preiner, M., Xavier, J. C., Zimorski, V. & Martin, W. F. The last universal common ancestor between ancient Earth chemistry and the onset of genetics. PLoS Genetics vol. 14 (2018).

27. Sofia, H. J. Radical SAM, a novel protein superfamily linking unresolved steps in familiar biosynthetic pathways with radical mechanisms: functional characterization using new analysis and information visualization methods. Nucleic Acids Res. 29, 1097–1106 (2001).

28. Ulrey, C. L., Liu, L., Andrews, L. G. & Tollefsbol, T. O. The impact of metabolism on DNA methylation. Hum. Mol. Genet. 14, 139–147 (2005).

29. Rodriguez-Granillo, A., Annavarapu, S., Zhang, L., Koder, R. L. & Nanda, V. Computational design of thermostabilizing D-amino acid substitutions. J. Am. Chem. Soc. 133, 18750–18759 (2011).

30. Annavarapu, S. & Nanda, V. Mirrors in the PDB: Left-handed-turns guide design with D-amino acids. BMC Struct. Biol. 9, 61 (2009).

