## Supplemental Figures/Table for "Alternating handedness motifs in proteins classify structure and cofactor binding"

S1

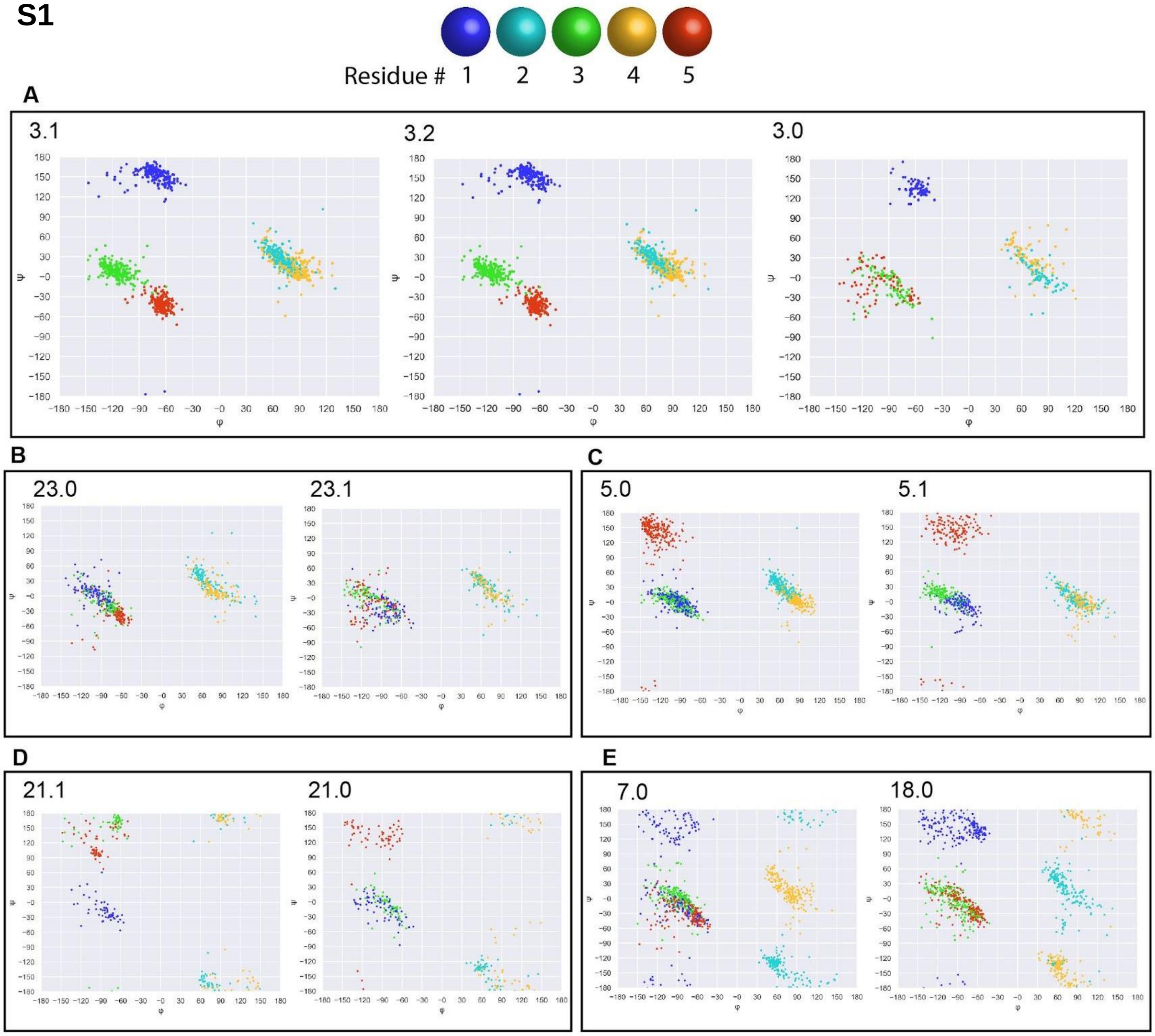


S2

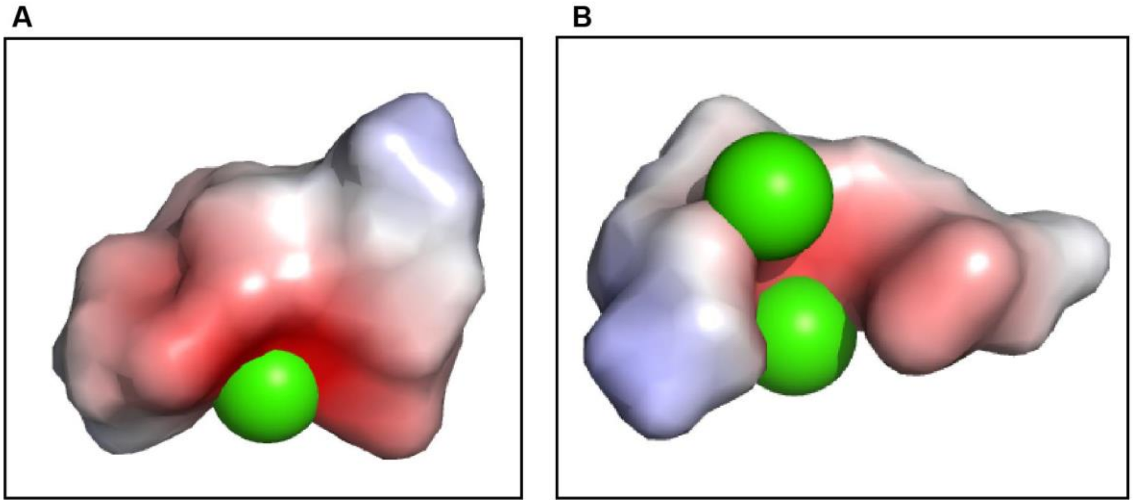

Table S1

| cluster | subcluster | # entries | avg curvature | std curve | consensus | percent hetatm | hetatm 1 | hetatm 2 | hetatm 3 |
|---|---|---|---|---|---|---|---|---|---|
| 3 | 1 | 213 | 90.10297049 | 8.848059675 | PRALARALAR | 0.840375587 | ADP, 43 | SO4, 37 | GDP, 20 |
| 3 | 2 | 222 | 85.76771369 | 12.96293969 | PRALARALAR | 0.653153153 | COA, 30 | ACO, 29 | SO4, 17 |
| 21 | 1 | 54 | 84.17513808 | 12.8297463 | ARPLPRNABR | 0.574074074 | CA, 33 | SO4, 1 | HEM, 1 |
| 5 | 0 | 217 | 101.6723388 | 12.11534375 | ARALARALBR | 0.571428571 | CA, 96 | NA, 7 | MG, 7 |
| 23 | 0 | 109 | 120.1263076 | 19.69498373 | ARALARALAR | 0.47706422 | SF4, 32 | F3S, 4 | SO4, 3 |
| 7 | 0 | 181 | 97.71897583 | 21.94045062 | ARPLARALAR | 0.215469613 | SAM, 7 | SAH, 7 | CL, 3 |
| 18 | 0 | 179 | 117.9874644 | 27.22083376 | PRALARPLAR | 0.212290503 | SAH, 7 | SO4, 6 | GOL, 3 |
| 17 | 0 | 180 | 90.64710005 | 19.00856402 | PRPLARALBR | 0.205555556 | ZN, 7 | FMN, 4 | MN, 4 |
| 23 | 1 | 91 | 100.5482729 | 23.49086312 | ARALARALAR | 0.197802198 | FES, 8 | K, 4 | SO4, 2 |
| 16 | 0 | 227 | 114.7432876 | 12.2514839 | ARALPRALAR | 0.145374449 | GDP, 7 | ACT, 3 | SO4, 2 |
| 6 | 0 | 239 | 61.40590197 | 15.85891308 | ARNAPRALPR | 0.133891213 | PLP, 6 | NA, 3 | SO4, 2 |
| 8 | 0 | 85 | 95.28614981 | 26.7163821 | BRPLARNABR | 0.129411765 | FMN, 2 | SO4, 1 | SAM, 1 |

| | | | | | | | | | |
|---|---|---|---|---|---|---|---|---|---|
| 20 | 0 | 145 | 101.0417595 | 16.06719629 | BRNABRPLAR | 0.124137931 | GOL, 4 | MG, 2 | TLA, 1 |
| 11 | 0 | 291 | 110.4656217 | 16.88079358 | PRALPRALBR | 0.12371134 | SAH, 5 | GOL, 4 | SO4, 3 |
| 13 | 0 | 165 | 118.0269211 | 23.68312939 | ARALBRPLAR | 0.121212121 | HEM, 7 | SO4, 2 | GOL, 2 |
| 9 | 0 | 173 | 104.1936857 | 16.28925659 | PRALBRPLAR | 0.115606936 | PO4, 2 | NDP, 2 | CMP, 2 |
| 1 | 0 | 174 | 108.9786433 | 15.9410156 | PRALPRALAR | 0.114942529 | SO4, 3 | GOL, 2 | FAD, 2 |
| 4 | 0 | 157 | 124.1339294 | 19.02866357 | ARALBRNABR | 0.114649682 | SO4, 3 | MG, 3 | CL, 2 |
| 21 | 0 | 62 | 82.42624199 | 23.09124958 | ARPLARNABR | 0.112903226 | NA, 2 | SAH, 1 | GOL, 1 |
| 3 | 0 | 74 | 102.9188911 | 19.0160215 | PRALARALAR | 0.108108108 | SO4, 3 | GOL, 2 | NAG, 1 |
| 15 | 0 | 199 | 64.54850955 | 17.4638038 | BRNAPRALPR | 0.105527638 | EOH, 6 | GOL, 2 | CA, 2 |
| 14 | 0 | 192 | 63.77822145 | 16.20316334 | BRNAPRALAR | 0.104166667 | CL, 4 | GOL, 2 | MG, 2 |
| 2 | 0 | 467 | 103.4730914 | 19.5191472 | PRALARALPR | 0.102783726 | SO4, 10 | CA, 10 | GOL, 8 |
| 5 | 1 | 148 | 72.69515745 | 16.63769158 | ARALARALBR | 0.101351351 | CA, 4 | MG, 3 | NA, 2 |
| 0 | 0 | 264 | 98.38445281 | 18.29927661 | PRALARPLPR | 0.098484848 | GOL, 4 | CL, 4 | SAH, 2 |
| 12 | 0 | 265 | 114.9046867 | 16.02899488 | ARALPRALPR | 0.098113208 | SO4, 8 | GOL, 3 | CA, 3 |
| 22 | 0 | 206 | 107.1814325 | 22.0738135 | ARALARNAPR | 0.097087379 | CA, 7 | SO4, 3 | SAH, 2 |
| 24 | 0 | 104 | 123.2171924 | 20.56687386 | PRALBRPLPR | 0.096153846 | MG, 2 | TRA, 1 | TLA, 1 |
| 19 | 0 | 228 | 110.9840695 | 24.90099397 | ARALARALPR | 0.065789474 | ZN, 3 | SF4, 1 | MLZ, 1 |
| 10 | 0 | 137 | 115.9980712 | 25.62770774 | ARALARPLAR | 0.065693431 | TRS, 1 | SAM, 1 | LHG, 1 |
